# Progressive degeneration in a new *Drosophila* model of Spinocerebellar Ataxia type 7

**DOI:** 10.1101/2023.11.07.566106

**Authors:** Alyson L. Sujkowski, Bedri Ranxhi, Matthew V. Prifti, Nadir Alam, Sokol V. Todi, Wei-Ling Tsou

## Abstract

Spinocerebellar ataxia type 7 (SCA7) is a progressive neurodegenerative disorder resulting from abnormal expansion of polyglutamine (polyQ) in its disease protein, ataxin-7 (ATXN7). ATXN7 is part of Spt-Ada-Gcn5 acetyltransferase (SAGA), an evolutionarily conserved transcriptional coactivation complex with critical roles in chromatin remodeling, cell signaling, neurodifferentiation, mitochondrial health and autophagy. SCA7 is dominantly inherited and characterized by genetic anticipation and high repeat-length instability. Patients with SCA7 experience progressive ataxia, atrophy, spasticity, and blindness. There is currently no cure for SCA7, and therapies are aimed at alleviating symptoms to increase quality of life. Here, we report novel *Drosophila* lines of SCA7 with polyQ repeats in wild-type and human disease patient range. We find that ATXN7 expression has age- and polyQ repeat length-dependent reduction in survival and retinal instability, concomitant with increased ATXN7 protein aggregation. These new lines will provide important insight on disease progression that can be used in the future to identify therapeutic targets for SCA7 patients.

## Introduction

Spinocerebellar ataxia type 7 (SCA7) is an inherited, autosomal dominant disease belonging to the family of polyglutamine (polyQ) neurodegenerative disorders^1,2^. The etiology of the polyQ family of diseases is in part, shared: toxic gain-of-function mutations—abnormal expansion of the triplet repeat CAG/A—yield longer than normal tracts of glutamine (Q)^3–8^. These elongated tracts cause protein misfolding and aggregation, particularly harmful to highly specialized neural cells. Although the expanded polyQ region is a shared feature in this neurodegenerative family, each causative disease protein is unique. Importantly, despite broad expression of disease genes, individual polyQ disorders affect different CNS regions and present distinct symptomology^9^.

Like other members of the polyQ family of disorders, SCA7 patients experience progressive neuronal loss in the cerebellum and brainstem that yield ataxia and early mortality, while unusually severe intergenerational repeat length instability, retinal degeneration and blindness are features that are unique to SCA7^10^. SCA7 results from abnormal CAG expansion in the *ATXN7* gene, whose protein product is part of Spt-Ada-Gcn5 acetyltransferase (SAGA), a highly evolutionarily conserved transcriptional coactivation complex^1,2^. SAGA has chromatin remodeling activities essential for RNA polymerase II transcription, and also modifies autophagy, cell-cell interactions, mitochondrial homeostasis and neuronal differentiation^10,11^. Additional work has implicated changes in gene expression, transcriptional regulation, and epigenetics in SCA7 pathogenesis, but the molecular mechanisms that underly its pathogenesis are not fully understood^12–15^.

Here, we report the generation and characterization of new *Drosophila* models of SCA7 that ectopically express full-length, Myc-tagged, human ataxin-7 (ATXN7) protein with normal (Q10) or expanded (Q92) polyQ, within patient range. Both Q10 and Q92 proteins had variable, tissue-specific toxicity, with 92Q conferring stronger progressive effects on retinal cell stability and longevity. The advantages of *Drosophila* models of SCA7 include their flexible and precisely controlled genetics, allowing for rapid, cost-effective, longitudinal investigations aimed at understanding pathogenesis and discovering therapeutic opportunities.

## Results

### Wild-type and polyQ-expanded models of SCA7

Techniques we successfully used in the past were used to generate new Gal4-UAS *Drosophila* lines that express Myc-tagged human ATXN7 with either Q10 or Q92^16–18^. Normal polyQ tracts range from 7-27 CAG repeats in *ATXN7* whereas full pathogenic penetrance occurs at 37-460 repeats^19^. A 5’ Myc tag was added to the full-length *ATXN7* gene with either a 10- or 92-triplet CAG repeat (Figure 1A) and sub-cloned into the pWALIUM10.moe vector (Figure 1B). This cloning strategy yields a single-copy, phiC31-dependent insertion into the attp2 site on the third *Drosophila* chromosome^20^, in the same integration site and the same orientation as our previously published polyQ family disease model flies^16–18^. We utilize this expression system and vector for its ability to easily modify protein expression, allowing us to both control the amount of disease protein expressed and compare across *Drosophila* polyQ models with similar protein expression levels^17,21,22^.

**Figure 1:**
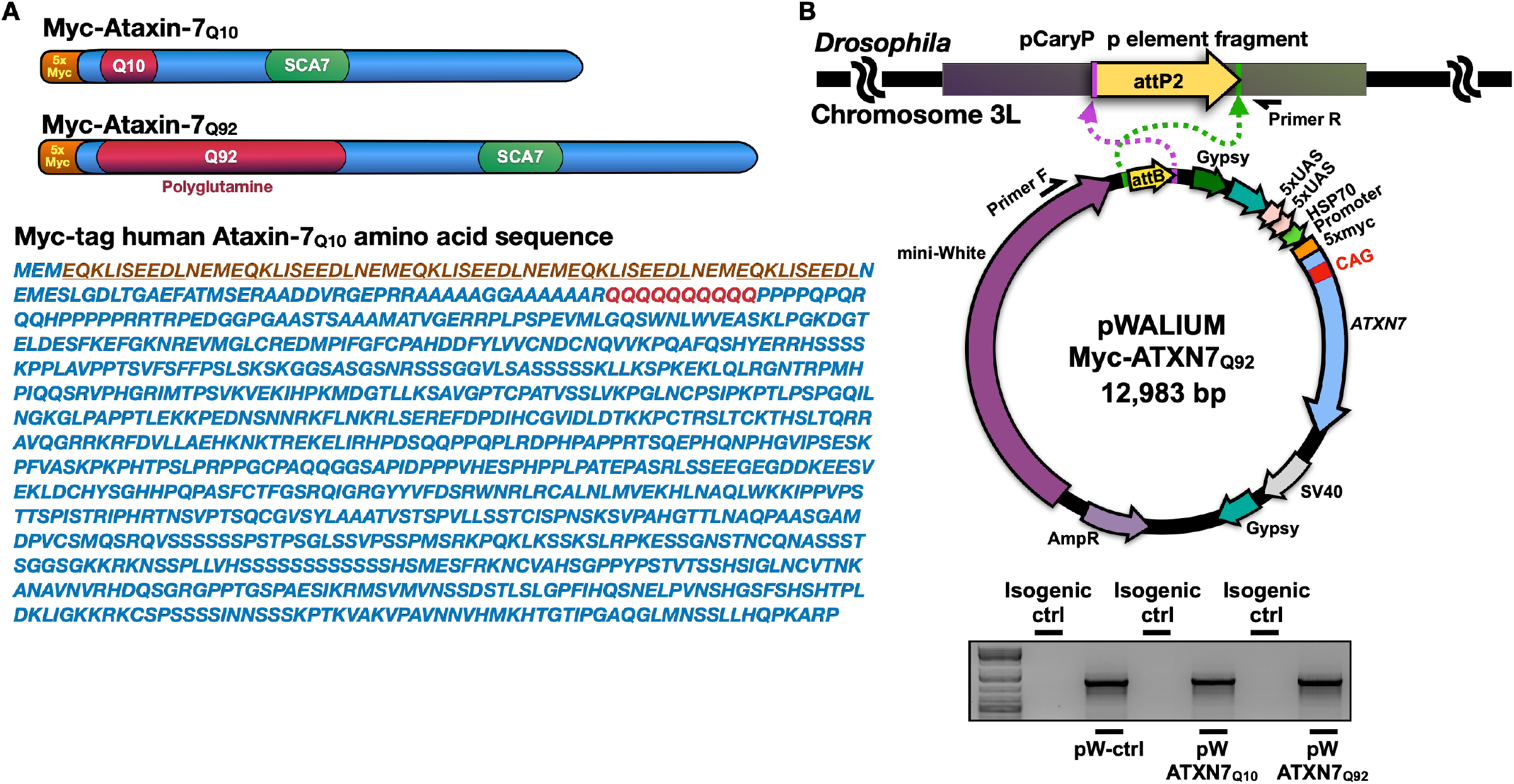
Generation of *Drosophila* SCA7 models. **(A)** Schematic of Myc-tagged ATXN7 Q10 and Q92 including construct-specific polyQ insertions. ATXN7 amino acid sequence, with 5X Myc tag (underlined) and polyQ repeat location (red) is also shown. **(B)** Top: diagrammatic representation of the cloning strategy used to insert ATXN7 cDNA into pWALIUM10.moe. Bottom: PCR reactions from genomic DNA indicating that the transgene was integrated into the correct site in the proper orientation.

### Expression of ataxin-7 in flies

We assessed ATXN7 protein and mRNA levels in one-day-old flies when UAS expression was driven in fly eyes (GMR-Gal4, Figure 2A), all neurons (elav-Gal4, Figure 2B), all glia (repo-Gal4, Figure 2C) or ubiquitously (sqh-Gal4, Figure 2D). When *ATXN7* expression was induced in either fly eyes (Figure 2A) or ubiquitously (Figure 2D) higher levels of Q92 mRNA and protein were observed in comparison to age-matched Q10 flies. ATXN7 mRNA and protein levels were similar, regardless of repeat length, when UAS expression was driven pan-neuronally (Figure 2B). We also examined ATXN7 mRNA and protein levels when UAS expression was confined to glial cells: although we observed lower levels of Q92 mRNA when compared to Q10, protein levels were higher with the Q92 variant (Figure 2C). Western blotting detected the presence of probable proteolysis products independent of driver (black arrows), suggesting high proteolytic activity of ATXN7 protein in all tissues tested, regardless of the polyQ length.

**Figure 2:**
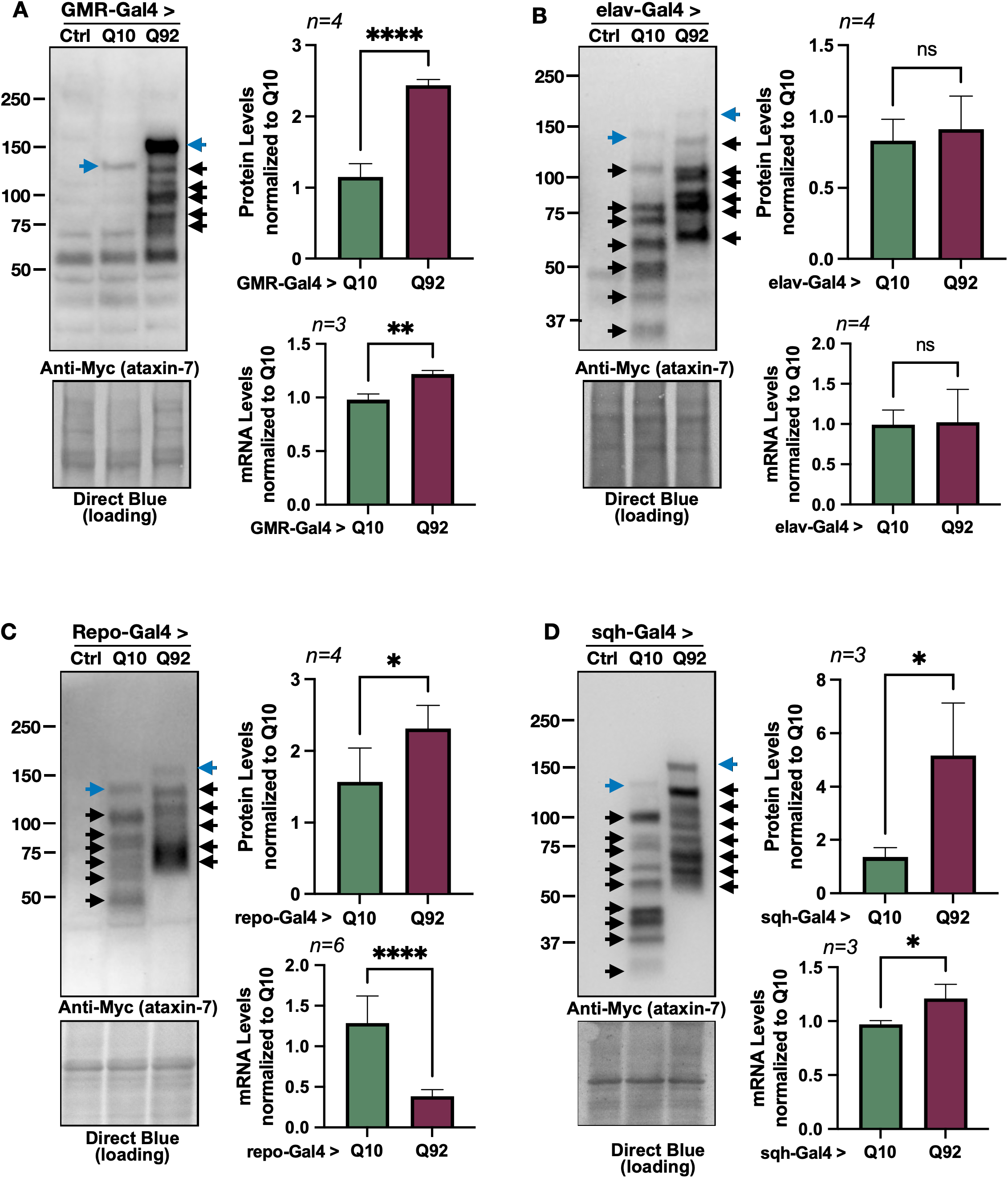
Expression of ataxin-7 in flies. Western blots from control, Q10, and Q92 flies in (**A)** fly eyes, (**B)** pan-neuronally, (**C)** in glia, and (**D)** ubiquitously. Quantifications are on the right of each blot. Q10 (105 kD) and Q92 (115 kD) indicated by blue arrows. Potential proteolysis products indicated with black arrows. The entire ataxin-7-positive signal was quantified in each case. qRT-PCR quantifications are found directly below Western blot quantifications. 10 fly heads (GMR-Gal4) or 5 whole flies (sqh-Gal4, elav-Gal4, repo-Gal4) were homogenized, depending on experiment. At least 3 biological replicates were used; ‘n’ indicated above panels. Significance determined by student t-test. *p>0.05, ****p<0.0001.

### Toxicity in SCA7 model flies

We have previously shown that ectopic expression of human, polyQ-expanded disease proteins reduces survival in multiple fly models of SCA neurodegeneration^16–18,23–28^. In order to determine any potential detrimental effects of ATXN7, we first tracked longevity in sex-separated, male and female flies expressing ATXN7 harboring either Q10 or Q92 in all tissues, or specifically in neurons, or glial cells (Figure 3).

**Figure 3:**
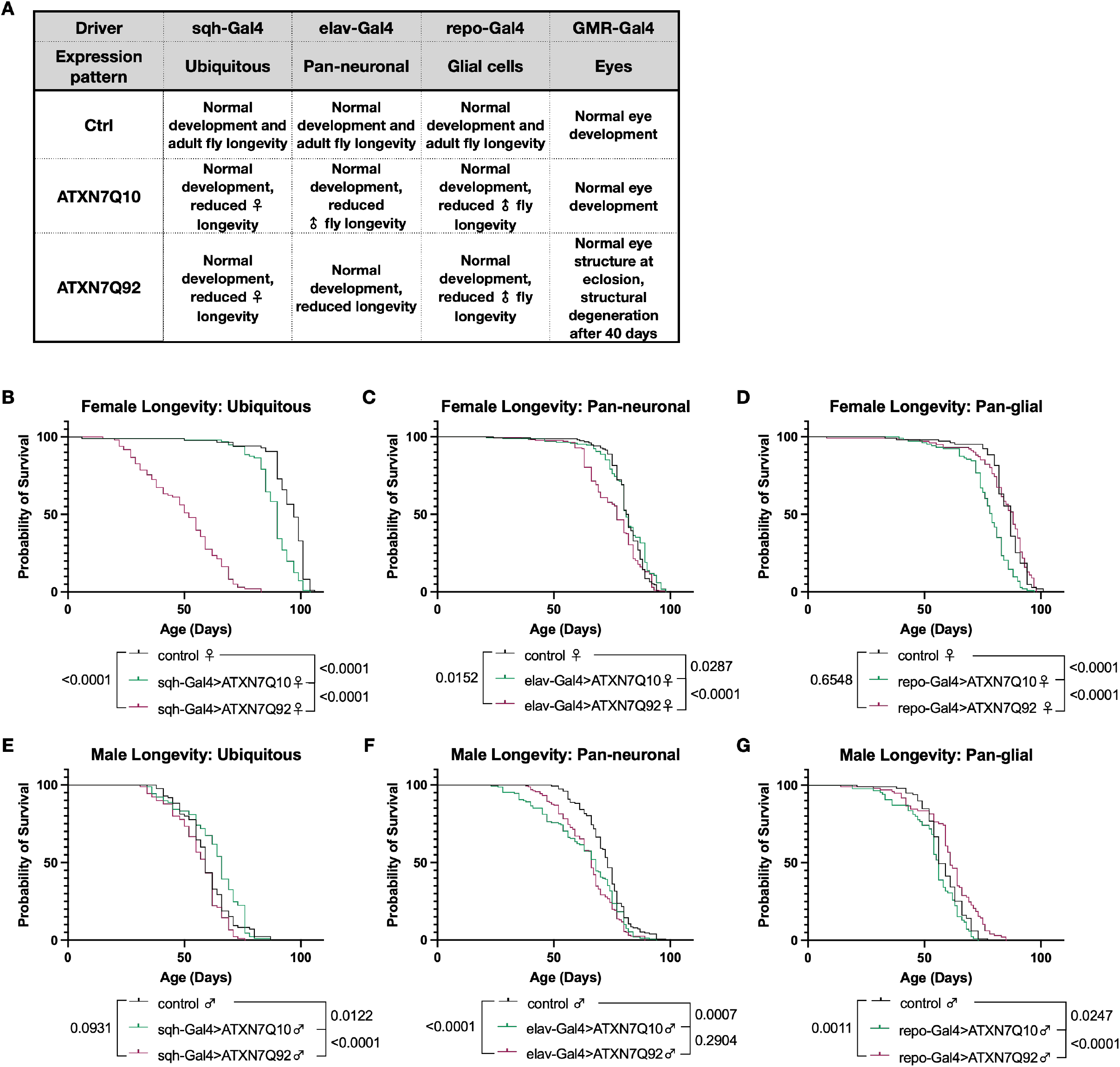
Effects of ataxin-7 expression on development and survival. A summary of experiments performed can be found in (**A).** Ubiquitous expression of Q10 reduced longevity in female flies but not males, and 92Q expression decreased female longevity **(B, E)**. Pan-neuronal expression of Q92 decreased female longevity **(C)** and both Q10 and Q92 male flies had reduced survival compared to controls **(F)**. Glial expression of ATXN7 Q10 reduced longevity in females and males, and glial Q92 expression did not **(D, G)**. n>200 flies for all groups, analyzed by log-rank.

Male and female controls flies developed normally and did not show reduced lifespan. Female flies ubiquitously expressing ATXN7 Q10 had shorter lifespan than background controls (heterozygous Gal4 flies backcrossed once into appropriate wild-type flies), and Q92 expression reduced lifespan further (Figure 3B). In male flies, ubiquitous expression of either Q10 or Q92 had less pronounced lifespan effects, although Q10 expression conferred modest longevity protection, particularly at mid-life (Figure 3E). Pan-neuronal expression of Q10 did not affect lifespan in female flies, but 92Q expression in female neurons decreased early-life longevity (Figure 3C). In contrast, male flies expressing ATXN7 Q10 or Q92 in all neural cells had shorter lifespan than age-matched controls (Figure 3F). When ATXN7 was restricted to glia, Q10 expression reduced longevity in both females and males, while Q92 expression was less toxic and did not negatively impact longevity (Figure 3D, 3G). Taken together, these results suggest that ATXN7 expression has sexually dimorphic, tissue-dependent effects on survival in *Drosophila*, and that expression of wild-type, human ATXN7 Q10 protein can have toxic effects in flies even in the absence of SCA7-associated polyQ expansions.

Expression of polyQ disease proteins in flies causes reduced motility, as we and others have shown before^16,27,29–41^. To test if ATXN7 expression negatively impacts mobility, we longitudinally measured 3 weeks of climbing speed in male and female flies expressing either Q10 or Q92 in the same cell populations as our longevity assessments; ubiquitously, neurons, and glia. In female flies, ubiquitous or glial expression of either Q10 or Q92 reduced climbing speed across ages (Figure 4A-C), and glial Q10 expression reduced climbing speed more than Q92 (Figure 4C). In contrast, neuronal ATXN7 expression did not negatively impact female climbing speed, and flies expressing Q10 in neurons had slightly better mobility than both control and Q92 flies (Figure 4B). In male flies, both Q10 and Q92 expression impaired mobility during the first week of adulthood whether ATXN7 expression was ubiquitous or specific to neurons or glia (Figure 4D-F). These results further support sexually dimorphic toxic effects of ATXN7 expression that are not fully dependent on polyQ repeat length.

**Figure 4:**
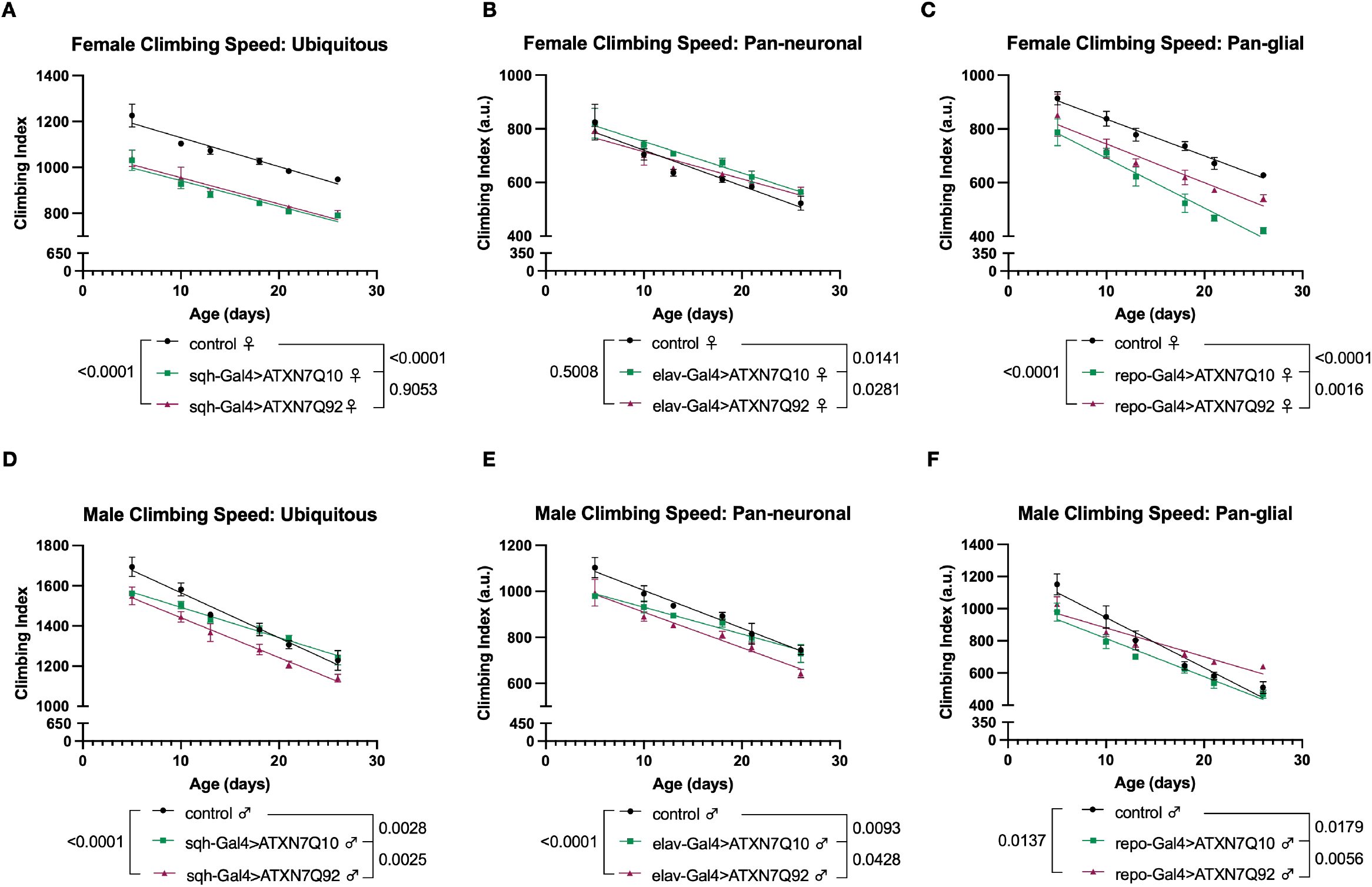
Effects of ataxin-7 expression on motility. **(A)** Both Q10 and Q92 expression reduced mobility in female flies. **(B)** Female flies expressing Q10 in neurons have slightly better climbing speed than background control flies. **(C)** Glial expression of Q92 reduces female climbing speed across ages, and Q10 expression impairs climbing further. **(D-F)** Male flies expressing Q10 or Q92 have lower climbing speed than age-matched controls at days 5 and 10 whether ATXN7 is expressed in all tissues **(D)**, neurons **(E)**, or glia **(F)**. n≥100 per genotype, per sex, analyzed by linear regression.

As noted above, glial expression of ATXN7 mRNA and protein levels were found in higher abundance than in age-matched Q10 flies (Figure 2C), a finding that correlates with a greater negative impact on longevity and mobility (Figure 3D, 3G, 4C, 4F). A similar length-dependent increase in ATXN7 protein was observed when expression was confined to fly eyes (Figure 2A), prompting us to examine whether differences in protein levels between Q10 and Q92 parallels toxicity in other tissues.

The *Drosophila* eye has been a stalwart model of neurogenerative diseases and genetic screens for many years^42–46^. We therefore used a fly eye specific SCA7 model to observe phenotypic neurodegeneration that may occur when ATXN7 is expressed, and to determine whether or not polyQ length or protein levels dictate degenerative phenotypes^16,47,48^. We have successfully used these tools in the past to detect changes that may be missed when disease protein expression is driven in other tissues^16,47,48^. When ATXN7 92Q was selectively expressed in fly eyes (GMR-Gal4), pseudopupil loss, an indicator of disorganization of underlying structures^16^, was present by adult day 56 (Figure 5A).

**Figure 5:**
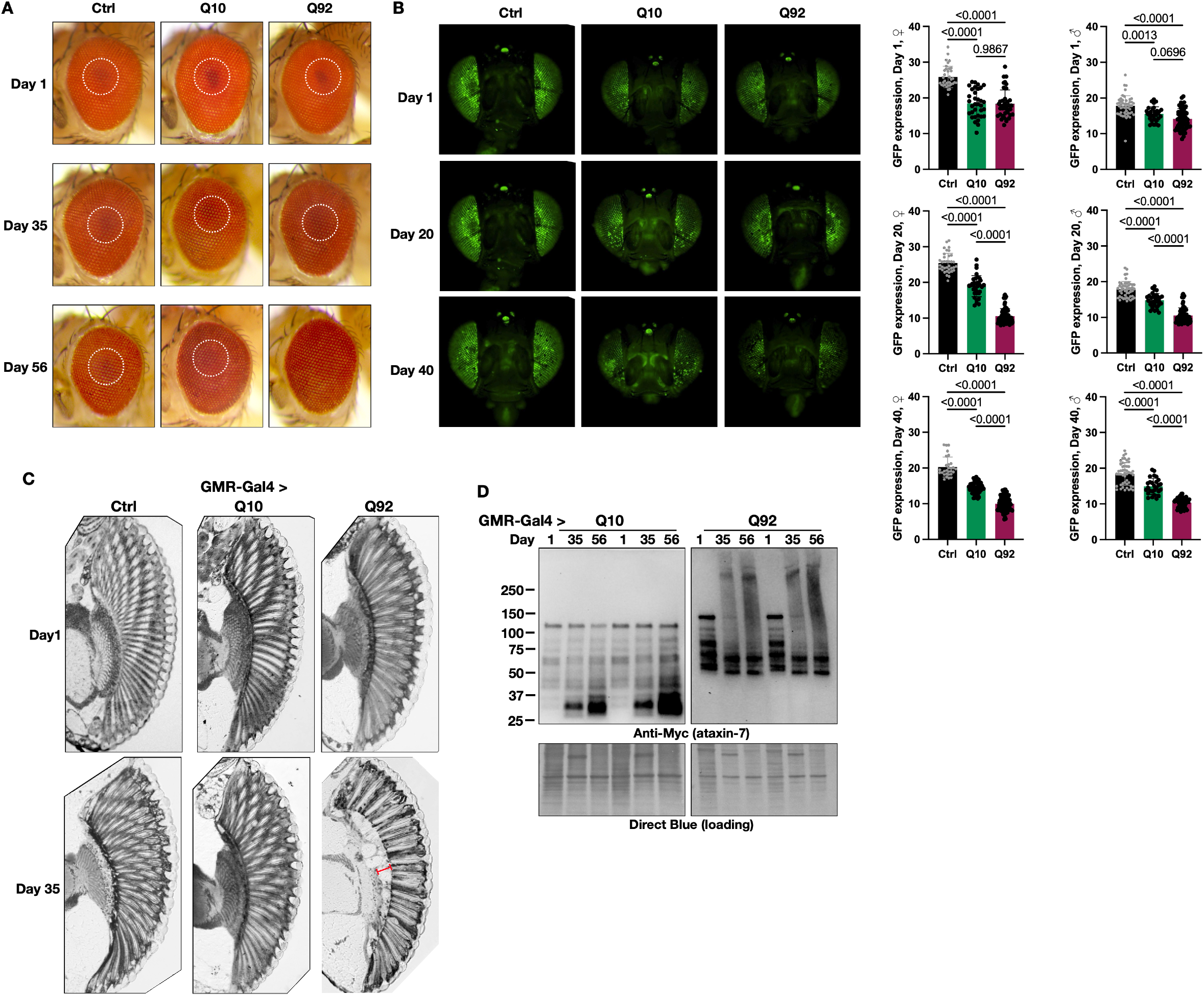
Effects of fly-eye specific ataxin-7 expression. **(A)** Pigmentation is normal in control and Q10 flies across ages. Q92 flies have pseudopupil (dotted circles) loss by adult day 56 n≥10. **(B)** CD8-GFP expression declines slightly with age in control flies (left panels, quantified at right; black histograms). Both Q10 (middle panels, green histograms) and Q92 (right panels, magenta histograms) have lower GFP fluorescence than age-matched control flies, and Q92 flies have lower GFP fluorescence than both control and Q10 flies by day 20 n≥29. **(C)** 5μM histological sections of control (left), Q10 (middle) and Q92 (right) flies at adult day 1 (upper panels). At adult day 56 (lower panels) Q92 flies have darkly stained aggregates, corneal disruption, loss of ommatidial boundaries (red bracket) and retinal disorganization. n≥6. **(D)** Western blots of ATXN7 in fly eyes over time. SDS-resistant species are present in the upper portions of the blot in Q92 flies by adult day 35. Lysates from 20 fly heads and at least 2 biological replicates performed per group.

To further dissect the timeline of retinal degeneration, we quantified fluorescent intensity of membrane-targeted GFP (CD8-GFP), an indirect measurement of retinal integrity^49^. In both male and female flies, GFP expression was already lower in one-day-old flies expressing either ATXN7 Q10 or Q92 when compared to background controls expressing CD8-GFP without ATXN7 (Figure 5B, top row and quantification on the right). By adult day 20, GFP expression was further reduced in flies expressing ATXN7 Q10, and 92Q expression in fly eyes exacerbated this phenotype, with more pronounced retinal cell loss observed by day 40 (Figure 5B, middle and bottom rows, respectively). These results suggest progressive, polyQ-length dependent retinal degeneration, detected earlier and with more sensitivity in this model than observation of external eye structure with light microscopy.

Histological examination of fly eyes supports these findings, revealing progressive degeneration that was more prominent with age and in Q92 model flies (Figure 5C). On adult day 1, eye morphology was unaffected regardless of genotype (Figure 5C, top row). In contrast, 92Q flies develop darkly stained aggregation products, loss of ommatidial boundaries (red bracket) and prominent retinal disorganization by adult day 35, suggesting age- and polyQ-length dependent neurodegeneration (Figure 5C, bottom row).

Lastly, we examined ATXN7 protein migration patterns in the fly eye-specific SCA7 model, a strong molecular indicator of polyQ family disease progression^9,50,51^. We have previously shown that aggregation of polyQ proteins precedes toxicity, and that aggregation levels of these proteins is an indicator of pathogenesis^16–18,23,26,48^. Eye-specific ATXN7 expression was visible in dissected Q10 and Q92 fly heads as early as adult day 1 (Figure 5D). SDS-resistant smears, indicative of aggregated species that migrate more slowly through SDS-PAGE gels, were apparent by day 35 and still present on day 56 in ATXN7 Q92 flies only. The SDS-resistant species were concomitant with loss of the primary SDS-soluble band of ATXN7 protein in the Q92 version (Figure 5D). Taken together, these results indicate progressive, polyQ-dependent toxicity that correlates with higher Q92 protein levels in these new SCA7 model flies, suggesting that ATXN7 toxicity coincides with both polyQ expansion and protein aggregation.

## Discussion

We described here two new *Drosophila* lines of SCA7 that rely on the expression of full-length, human ATXN7 protein with a polyQ repeat length in either wild-type (Q10) or human SCA7 disease range (Q92). Compared to control flies, both Q10 and Q92 expression in neurons, fly eyes, or ubiquitously had variable, progressive degenerative effects; Q92 expression was particularly deleterious to survival and in fly eyes, concomitant with increased aggregation over time in the latter tissue. Our initial characterization of these new models opens the door for the generation of new isogenic lines that precisely dissect the molecular mechanisms driving disease progression in SCA7.

Fly eyes have been a valuable tool to aid our understanding of proteotoxic neurodegenerative disease progression^17,18,28,49,52–56^. In this report, we observe ATXN7-dependent eye degeneration in an age- and polyQ length-dependent manner. This finding is especially important in the context of SCA7, as progressive blindness is a clinical presentation that is unique to this disorder^6,19^. SCA7 is broadly classified into three categories: adult-, young-adult, and early-childhood—classifications based on clinical presentation, age of onset, and polyQ repeat length^57^. Although common in earlier-onset diseases, patients who present in the fifth decade of life and later do not typically experience visual impairment^19,57^. With our fly models, we observe age- and polyQ repeat length-dependent decreases in GFP fluorescence when Q10 or Q92 proteins are expressed in the fly eye; histological sections reveal severe and progressive retinal deterioration in Q92 model flies, without clear impact from the Q10 counterpart. These new tools will provide a valuable platform by which to determine the timeline of retinal phenotypes as well as potential genetic or pharmacological modifiers of blindness in SCA7 patients.

In contrast, when ATXN7 is expressed in all tissues, neurons, or glial cells, toxic effects on survival and motility are sex- and tissue-specific and do not fully depend on polyQ repeat length. Furthermore, we report a correlation between protein levels and toxic phenotypes, particularly when ATXN7 was expressed in fly eyes or ubiquitously. We and others have observed detrimental effects from overexpression of wild-type, human polyQ proteins in *Drosophila*^17,58^; the mechanisms underlying their pathogenicity are unclear. In *Drosophila,* loss of endogenous *Atxn7* yields neurodegenerative phenotypes that are similar to overexpression of polyQ-expanded human ATXN7^59,60^. *Atxn7* mutation uncouples SAGA’s normal function as a transcriptional coactivator from its protein quality control function, resulting in aberrant gene expression that yields reduced lifespan, impaired motility, and retinal degeneration^59–61^. It is possible that toxicity from Q10 expression results from disrupted endogenous Atxn7 function, while both loss of function and aggregation-related gain-of-function contribute to Q92 toxicity. Using these new lines, future work may differentiate the relative contribution of protein level, aggregation, and loss of endogenous ATXN7 function to SCA7 pathogenesis.

We report here sex-specific effects of ATXN7 Q10 and Q92 expression on both survival and motility. Clinically, SCA7 is dominantly inherited and does not present in a gender-specific way^62^. This difference between humans and our fly models of SCA7 can be leveraged toward the identification of therapeutic targets, looking for unique, specific factors that are protective. Sex determination in *Drosophila* is cell autonomous and reversible, controlled through alternative splicing of a single gene^63^. We and others have taken advantage of this genetic flexibility to identify the cellular source of variation between male and female flies^63–70^. As described above, male SCA7 model flies are protected from early mortality conferred by ubiquitous ATXN7 expression, but more sensitive to neurodegeneration when expression is confined to neurons. Follow-up investigations could be geared toward identifying molecular targets that are protective in particular sexes, and in particular tissues, with the goal of modulating those pathways to prevent neurodegeneration across sexes and age.

Finally, examination of protein levels in our Q10 and Q92 flies indicates the presence of proteolytic fragments that differ depending on age, polyQ length, and the tissue in which they are expressed (see Figures 2, 5D). Proteolytic fragmentation of disease proteins is not uncommon in polyQ family disease models and is thought to modify aggregation, function, localization, and toxicity of disease proteins^71–75^. In SCA7, toxic protein fragments cause increased nuclear retention of ATXN7 and impair its normal functional interaction with SAGA^71^, and may yield post-translational modifications that prevent normal degradation^76^. Our results support these findings, particularly in our fly eye specific SCA7 model. Here, we observe gradually accumulating proteolytic cleavage products of Q10 expression, whereas Q92 fragments shift over time, a finding that correlates with more severe retinal degeneration. These models can be leveraged to precisely identify how proteolytic cleavage and posttranslational modifications define aggregation susceptibility and proteotoxicity in SCA7.

Together, these new *Drosophila* models provide a versatile, cost-effective system under precise genetic control that will enable expedient and targeted examination of the molecular pathways that drive SCA7 neurodegeneration. These tools will be used in the future to guide the generation of new therapeutics for SCA7 patients and their families.

## Methods

### Fly Stocks and Maintenance

elav-Gal4 (#458) and GMR-Gal4 (#8121) were obtained from the Bloomington *Drosophila* Stock Center. w-; UAS-CD8-GFP; GMR-Gal4 was described previously^49^. Gifted stocks used in this study were sqh-Gal4 (Daniel Kiehart, Duke University), repo-Gal4 (Vanessa Auld, University of British Columbia), y, w; +; attP2 (Jamie Roebuck, Duke University) and w^1118^ (Russ Finley, Wayne State University). To generate transgenic lines that express ATXN7Q10 (Q10) and ATXN7Q92 (Q92) through the Gal4-UAS system^77^, the DNA sequence of full length, human ATXN7 was inserted into pWALIUM10-moe plasmid (DNA Resource Core at Harvard Medical School, MA, USA) with a 5’ Myc tag and a construct-specific (10Q or 92Q) polyQ expansion. Constructs were injected by the Duke University Model System Injection Service into y, w; +; attP2. For transgene verification, genomic DNA were extracted from different founder lines using DNAzol (ThermoFisher, Waltham, MA USA) and PCR-amplified using the below primers.

white-end-F: 5′-TTCAATGATATCCAGTGCAGTAAAA-3′

attP2-3L-R: 5′-CTCTTTGCAAGGCATTACATCTG-3′.

Flies were housed in a 25^°^C incubator on a 12:12h light:dark cycle at 40% relative humidity. Control flies for all Gal4-UAS experiments consisted of heterozygous Gal4 lines backcrossed into y, w; +; attP2 and/or w^1118^ flies, depending on experiment. Adult progeny were synchronized by collecting within 12 hours of eclosion over a 24 hour time period. Groups of 10-20 age- and sex-matched flies were immediately transferred into narrow polypropylene vials containing 5mL of standard 2% agar, 10% sucrose, 10% yeast with appropriate preservatives. Food vials were changed every second to third day.

### Western Blots

Five whole flies or twenty fly heads per biological replicate, depending on experiment, were homogenized in boiling lysis buffer (50 mM Tris pH 6.8, 2% SDS, 10% glycerol, 100 mM dithiothreitol), sonicated, boiled for 10 min, and centrifuged at 13,300xg at room temperature for 10 min. Western blots were developed ChemiDoc (Bio-Rad, Hercules, CA, USA) and quantified with ImageLab (Bio-Rad, Hercules, CA, USA). For ATXN7, protein levels were measured by quantifying the entire lane (Anti-Myc). Q92 expression was normalized to Q10 expression, taking into account total protein for each genotype using direct blue. Primary antibody used was Mouse anti-Myc (9B11) (1:1000, Cell signaling, Danvers, MA, USA); Secondary antibody: peroxidase-conjugated anti-mouse (1:5000, Jackson Immunoresearch, West Grove, PA, USA). For direct blue staining, PVDF membranes were submerged for 10 min in 0.008% Direct Blue 71 (Sigma-Aldrich, St. Louis, MO, USA) in 40% ethanol and 10% acetic acid, rinsed in 40% ethanol/10% acetic acid, air dried, and imaged. Western blots were performed using at least 3 biological replicates, and statistical analysis (student t-test) was performed in GraphPad Prism (San Diego, CA, USA).

### qRT-PCR

Total RNA was extracted by using TRIzol reagent (Life Technologies, Waltham, MA, USA). Extracted RNA was treated with TURBO DNAse (Ambion, Waltham, MA, USA) to remove contaminating DNA. Reverse transcription was performed with the High-Capacity cDNA Reverse Transcription Kit (ABI, New York, NY, USA). Messenger RNA levels were quantified by using the StepOnePlus Real-Time PCR System with Fast SYBR Green Master Mix (ABI). rp49 was used as internal control. At least three biological replicates were used, and statistical analysis (student t-test) was performed in GraphPad Prism (San Diego, CA, USA).

Primers:

Atxn7-F: 5′- CTGCTCTCATCTACCTGCATCTC-3′;

Atxn7-R: 5′- TAGTGCTGTTACCAGAAGACTCCTT-3′;

rp49-F: 5′- AGATCGTGAAGAAGCGCACCAAG-3′;

rp49-R: 5′- CACCAGGAACTTCTTGAATCCGG-3′.

### Lifespan

At least 200 adults were age-matched and separated by sex within 12 hours of eclosion. Flies were transferred to narrow polypropylene vials containing 5mL of standard 2% agar, 10% sucrose, 10% yeast food. Flies were transferred and scored for death events every 2-3 days until no flies remained. Survival curves were analyzed by log-rank in GraphPad Prism (San Diego, CA, USA).

### Climbing speed

Negative geotaxis was assessed using a modified Rapid Iterative Negative Geotaxis (RING) assay in groups of at least 100 flies as described^27,41,78^. Briefly, vials of 20 flies were briskly tapped down, then measured for climbing distance after 2s of inducing the negative geotaxis instinct. For each group of vials, an average of 5 consecutive trials was calculated and batch processed using ImageJ (Bethesda, MD). Flies were longitudinally tested 2-3 times per week for 3 weeks. Between assessments, flies were returned to food vials and housed normally as described above. Negative geotaxis results were analyzed using linear regression (slope, y-intercept) in GraphPad Prism (San Diego, CA, USA). All negative geotaxis experiments were performed in duplicate, with one complete trial shown in each graph.

### Fluorescence measurements

All fluorescence images were taken with an Olympus BX53 microscope and CellSens software (Olympus, Waltham, MA, USA). The same objective (4X) and camera settings (ISO 200, exposure time 500 ms capture time for each sample) were used for all images. Control and experimental flies were all imaged on the same day. ImageJ’s freehand tool was used to outline and to quantify fluorescence readings from each fly eye. GFP expression was statistically analyzed by ANOVA in GraphPad Prism (San Diego, CA, USA). n≥29 flies for all groups.

### Histology

Adult flies whose proboscises and wings were removed were fixed in 2% glutaraldehyde/2% paraformaldehyde and 0.05% Triton X-100 in phosphate buffered saline overnight at 4°C. Fixed flies were later dehydrated in a series of 30, 50, 75 and 100% ethanol and propylene oxide, embedded in Poly/Bed812 (Polysciences, Warrington, PA, USA), sectioned at 5 μm and then stained with Toluidine Blue. n≥6 flies for all groups.

## Acknowledgements

We extend our gratitude to Pegi Laci and Racquel Harrison for help with conducting lifespan and mobility assays.

## Author contributions

AS: conceptualization, data curation, software, formal analysis, validation, investigation, visualization, methodology, and writing and editing.

BR: data curation, validation, investigation, methodology.

MVP: data curation, validation, investigation, methodology.

NA: data curation, validation, investigation, methodology.

KL: data curation, validation, investigation, methodology.

SVT: conceptualization, data curation, funding acquisition, software, formal analysis, validation, investigation, visualization, methodology, and writing and editing.

W-LT: conceptualization, data curation, software, formal analysis, validation, investigation, visualization, methodology, and writing and editing.

## Data Availability Statement

Lines, plasmids and source data are available upon request. The authors affirm that all data necessary for confirming the conclusions of the article are present within the article and figures.

## Competing Interests Statement

The authors declare that they do not have any conflicts of interest to disclose.

## Notes

Grant information: This study was supported in part by NINDS R01NS086778 (SVT).

### Competing Interest Statement

The authors have declared no competing interest.

